# Scaling up tree diversity inventories across Amazonian ecosystems using field spectroscopy

**DOI:** 10.1101/2025.03.26.645444

**Authors:** Hilana Louise Hadlich, Jochen Schöngart, Florian Wittmann, Caroline C. Vasconcelos, Caroline L. Mallmann, Maíra L. G. Conde, Priscila Amaral de Sá, Layon O. Demarchi, Gisele Biem Mori, Maria T. F. Piedade, Flavia Machado Durgante

## Abstract

- Species identification in Amazonian forest inventories is challenging due to a shortage of taxonomists, high biodiversity, and morphological similarities leading to taxonomic errors. Near-infrared spectroscopy (NIRS) is a promising tool for improving species identification efficiency and reliability.
- This study assessed the effectiveness of NIRS in discriminating against 26 abundant tree species across three Amazonian ecosystems: upland forest, white-sand ecosystems, and floodplain forest, using spectral data from different tree tissues—outer bark, inner bark, and fresh leaves. Each tissue was tested using Linear Discriminant Analysis (LDA) spectral models with two cross-validation methods: leave-one-out and 70/30 hold-out.
- Results showed high discrimination accuracy for all tissues and ecosystems. The general models achieved 86% accuracy for outer bark, 97% for inner bark and 98% for fresh leaves. The most informative spectral bands varied by tissue type: SWIR I (1300–1900 nm) for outer bark, and VIS (400–700 nm) + SWIR I (1300– 1900 nm) for inner bark and fresh leaves. A general model integrating species across ecosystems confirmed NIRS as an effective tool for in-field tree identification. These findings highlight the potential of VIS-NIR spectroscopy to Amazonian biodiversity inventories, contributing to more accurate species identification, refining forest management and conservation efforts.

## Introduction

Conserving biodiversity of tropical forests is a challenge, especially given the lack of knowledge about the totality of species that occur in the vast mosaic of habitats and ecosystems that compose these forests. Although there have been many efforts to find out how many tree species exist in the Amazon (ter Steege *et al*., 2013, 2016, 2019; Cardoso *et al*., 2017; Oliveira-Filho *et al*., 2021), and it is estimated that there are more than 15,000 tree species (ter Steege *et al*., 2020), but only around 6,000 have been cataloged (Cardoso *et al*., 2017). The challenges of delimiting and quantifying species persist due to taxonomic uncertainties (Draper *et al*., 2020; Stropp *et al*., 2022) and lack of botanical collections in remote regions (Nelson *et al*., 1990; Hopkins, 2007, 2019). This uncertainty directly affects biodiversity inventories that monitor tropical forests and compromises subsequent activities that depend on this information, such as forest management and conservation, as well as scientific research and ecological modeling. Although the problem of species identification in the Amazon is well-known in forest management (Procópio & Secco, 2008; Cysneiros *et al*., 2018), it is still neglected due to the lack of efficient and quick inspection techniques. The consequences of misidentification can be both economic and ecological, including the overestimation of commercially valuable species populations, the compromise forest product transaction integrity (Baraloto *et al*., 2007, Procópio & Secco, 2008; Gaui *et al*., 2019), and facilitation of rare species exploitation and illegal logging. Therefore, mitigating identification errors in forest inventories is an urgent priority. It is essential to focus on training new parataxonomists and developing innovative technologies and methodologies to optimize the process.

Field monitoring of the Amazon rainforest is conducted through forest inventories, where the first step is to ensure species identification. This is a slow, laborious process and requires a high level of taxonomic knowledge. Identification begins in the field facing difficulties due to the high morphological similarity between tropical species, making this activity subjective and inaccurate (Gomes *et al*., 2013) and the scarcity of botanical taxonomists, since the samples collected in the field must be validated with reference collections in herbarium. However, the absence of reproductive material for accurate identification can lead to the accumulation of errors in herbarium, perpetuating misidentifications. This issue is exacerbated by a mismatch of over 50% in the names of tropical species (Hopkins, 2007, Procópio and Secco, 2008; Gomes *et al*., 2013; Goodwin *et al*., 2015). Therefore, those responsible for forest monitoring need species verification techniques beyond field identification to ensure quality control and reduce the errors associated with identification.

Some approaches have been developed to solve the taxonomic challenges, such as DNA barcoding, mass spectroscopy, near-infrared spectroscopy (NIRS) in the field and the laboratory, and imaging spectroscopy. Each of these techniques has its advantages and limitations, and their integrated use is recommended to minimize identification errors (Prata *et al*., 2018; Damasco *et al*., 2019; Draper *et al*., 2020). Among the available tools, the *in situ* optical spectroscopy technique is a way to evaluate biodiversity and support forest management with the automation of data collection (Juola *et al*., 2023) and is suitable to optimize forest inventories (Hadlich *et al*., 2018). NIRS uses light radiation on organic samples to generate reflectance or absorbance spectra in the near-infrared regions. Multivariate techniques can characterize the sample quantitatively and qualitatively (Foley *et al*., 1998; Pasquini, 2003, 2018; Tsuchikawa, 2007). The spectral response depends on the sample’s chemical composition, cellular structure, and internal morphology (Ponzoni, 2002), and each species is assigned a spectral signature.

Over the years, NIRS methodologies have been developed to discriminate between species, employing various data sources and both laboratory and field equipment. The technique has already been tested on samples of herborized leaves (Durgante *et al*., 2013; Lang *et al*., 2015; Botelho, 2017; Meireles *et al*., 2020); laboratory-dried woods (Pigozzo, 2011; Braga *et al*., 2011; Pastore *et al*., 2011, Bergo *et al*., 2016; Eugenio-da-Silva *et al*., 2024), fresh leaves in field (Asner *et al*., 2014; Mallmann *et al*., 2023) and bark of standing trees (Hadlich *et al*., 2018; Juola *et al*., 2022; Juola *et al*., 2023), demonstrating remarkable potential for these applications. In an integrative context, NIRS has also been effective in assisting the delimitation of species with cryptic morphological variation (Prata *et al*., 2018; Damasco *et al*., 2019), even hypothesizing new species (Vasconcelos *et al*., 2020, 2021, in press), which shows the importance of using multiple tools to solve eventual taxonomic errors, especially in the highly diverse Amazonian flora.

The application of NIRS to improve species identification has shown promising results in the field mainly in upland forests based on bark (Hadlich *et al*., 2018) and leaf spectra (Asner *et al*., 2014, Mallmann et al. 2023). However, its applicability in different Amazonian ecosystems remains largely unexplored. The Amazon is made up of distinct ecosystems in terms of floristics, soil types, topography and water availability, which form mosaics of landscapes, each with specificities that must be considered when monitoring biodiversity. The predominant vegetation is non-flooded upland forest (also known as *terra-firme* forest), which covers about 70% of the basin. The remaining 30% consists predominantly of wetland complexes, most of them forested, and which experience seasonal flooding (Junk *et al.,* 2011, 2014). Additionally, other ecosystems are interspersed throughout the biome, including savannas, coastal vegetation, montane forests, and the white-sand ecosystem (locally known as *campinarana*), the latter covering approximately 5% of the basin area (Adeney *et al*., 2016). Given this environmental heterogeneity, NIRS-based identifications must be tailored to the specific conditions of each ecosystem to improve the accuracy of species identification as the spectral variation between tree species is influenced by chemistry and environmental conditions (Clark & Roberts, 2012).

Inventorying Amazonian ecosystems is a complex task that requires strategic planning to minimize cost and maximize efficiency. Considering the specific limitations of each ecosystem during the planning phase can significantly optimize the process. To improve forest inventories using visible and near-infrared spectroscopy for in-field species identification, three key questions have been addressed in this study: How effective are different plant tissues in predicting species identification across different Amazonian ecosystems? Is this efficiency consistent for all ecosystems together? Which wavelengths are the most informative for species recognizing across different tissues? The findings will provide a practical framework for improving inventory methodologies, ensuring greater accuracy while reducing costs. This research will contribute to more reliable biodiversity assessments and better-informed conservation and management strategies by shifting from uncertain identifications to a high-precision approach.

## Materials and Methods

### Study sites

Spectral data were systematically collected from established permanent plots as part of the Long-term Ecological Research Program (PELD) of the Working |Group Ecology, Monitoring and Sustainable Use of Wetlands (MAUA) within the Amazon Tall Tower Observatory (ATTO site). The ATTO site is situated in the Uatumã Sustainable Development Reserve (02°10’30’’ S; 59°00’30’’ W), approximately 150 km northeast of Manaus, Amazonas, Brazil (Fig. 1). The climate within the study area is the equatorial pluvial (Af) type, following the Köppen-Geiger classification system (Alvares *et al*., 2013). The region experiences an average annual rainfall of approximately 2,030 mm and an average temperature of 28 °C. The dry season extends from June to October and is characterized by reduced rainfall, generally below 100 mm (Andreae *et al*., 2015).

**Fig. 1.**
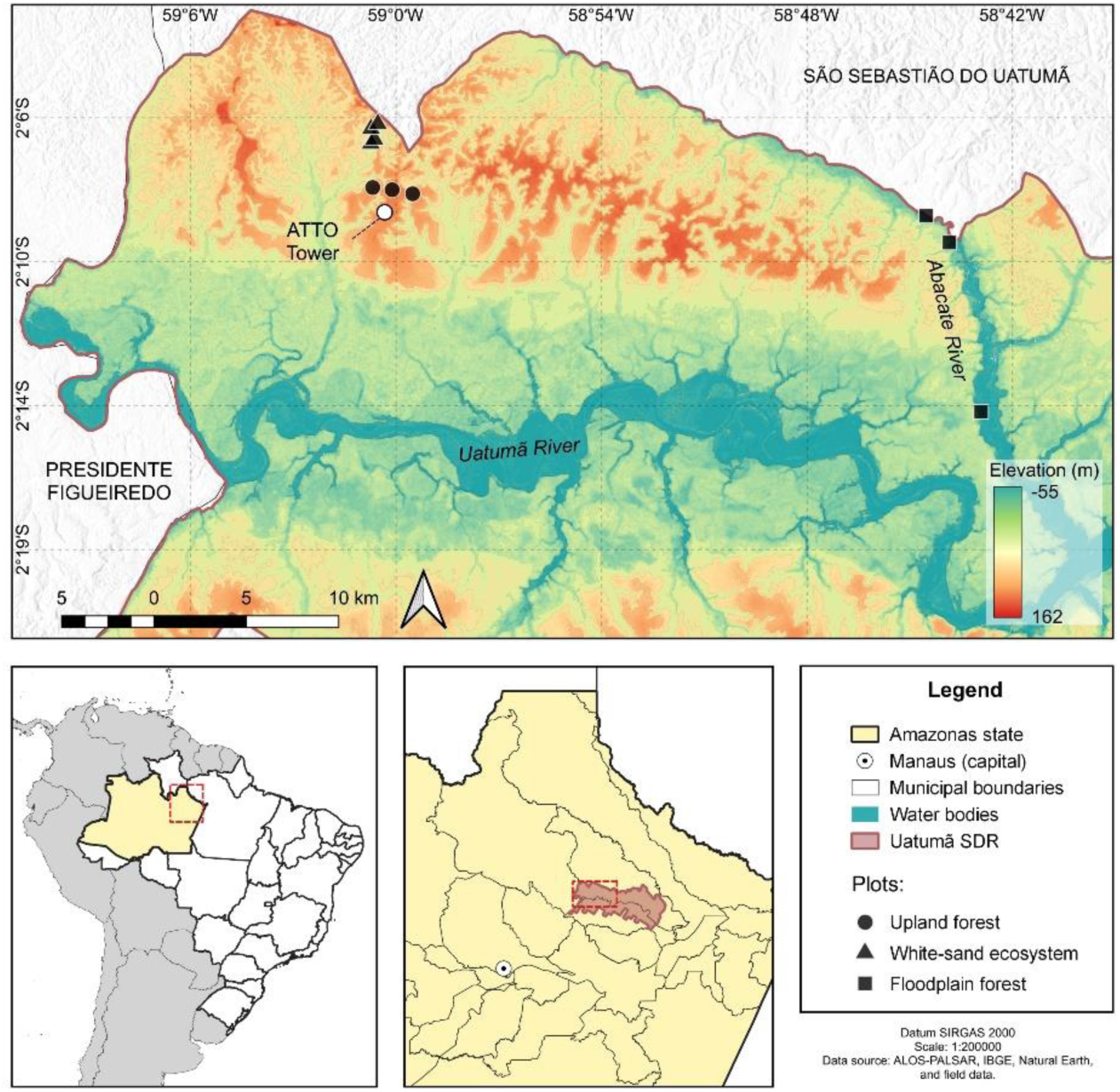
Map of the study area in Central Amazon, showing the sampling permanent plots where spectral data were collected from tree species. Solid black markers represent plots in upland (circles), white-sand (triangles), and floodplain forests (squares).

The present research was developed in three distinct forest ecosystems: upland forest (*terra-firme*), white-sand ecosystem (*campinarana*), and floodplain forest (*igapó*).

The upland plots are located on plateaus near the ATTO tower (Fig. 1), reaching a maximum altitude of approximately 130 m. The upland forest grows on clayey soil, Ferralsols (85.3% clay), which have good moisture retention capacity (Chauvel *et al*., 1987; Andreae *et al.,* 2015). It is a dense forest, with a mean height of 21 m and canopy trees reaching up to 45 m. The studied plots comprise 54 botanical families, with Fabaceae, Lecythidaceae, Sapotaceae, Burseraceae, and Euphorbiaceae being the most abundant. Based on previous forest inventory data, a total of 179 tree species were recorded, with *Protium apiculatum* and *Pouteria* sp. as the most abundant species (Andreae *et al.,* 2015).

The white-sand ecosystem of the ATTO site is positioned between the ancient river terraces and the plateau slopes (Targhetta *et al*., 2015). White-sand ecosystems are characterized by nutrient-poor, highly acidic, and easily leached soils. The soil is predominantly sandy, with a sand fraction of 93.4 ± 1.5% (Targhetta *et al*., 2015), resulting in low water retention capacity. According to Demarchi *et al*. (2022), the white-sand ecosystem of the RDSU are composed of six distinct phytophysiognomies, varying from open to dense white-sand ecosystem, in which the dense phytophysiognomie are differentiated by the reach of the water table to the surface. 167 taxonomically identified species were recorded in the ATTO site, 30% of which are endemic. In this study, four plots were selected of the open to intermediate white-sand ecosystem (Fig. 1) that suffer the most significant water deficit, especially in the dry season. The mean height of the trees in this environment is between 15-20m, with some emergent trees reaching 25-30m high in the canopy. In these plots, 40 families were recorded, with Fabaceae, Humiriaceae, Sapotaceae, Chrysobalanaceae, and Rubiaceae being the most abundant. There were 146 species, of which *Sacoglottis guianensis* was the most abundant, followed by *Aldina heterophylla* and *Pradosia schomburgkiana*, both specialists in Amazonian white-sand ecosystem (Demarchi *et al*., 2022).

The floodplain forest plots are located along the Abacate River, the main tributary of the Uatumã River (Fig. 1). This river is classified as a blackwater (locally known as igapó) that originate in the Guiana Shield and the Central Brazilian Shield (Junk *et al*., 2011). These waters are poor in suspended sediments and carry few nutrients, so the soils associated with blackwater rivers are of low fertility and high acidity (Junk *et al*., 2015).

The permanent plots used were those of the low and medium igapó of the Abacate River. These floodplain forests remain preserved, as their hydrological cycle has not been impacted by the Balbina Dam (Schöngart *et al*., 2021). The flooding regime varies from 1 to 3 m for 50 to 100 days a year in medium igapó. The species sampled from the low igapó such as *E. tenuifolia*, *M. acaciifolium*, *H. brasiliensis*, *M. tamaquarina* (Table 1) experience flooding from 96 to 127 days a year and can reach up to 7 meters in height (Lobo *et al*., 2019, Householder *et al*., 2021). The percentages of sand and clay are 57.0% and 27.4%, respectively (Mori, 2019). Therefore, the soil is characterized as clay-sandy. The trees in these environments have a mean height of 13.5 m and can reach a height of approximately 30 m. The flora of the plots is represented by 26 botanical families, with Lecythidaceae, Ochnaceae, Sapotaceae, and Fabaceae having the most abundance. A total of 65 species were recorded, with *Elvasia calophyllea*, *Eschweilera albiflora*, *Pouteria pachyphylla*, and *Eschweilera tenuifolia* as the most abundant species (Schöngart *et al*., 2021).

**Table 1.** Selected tree species in each ecosystem. A minimum of ten individuals were used for each species.

| Upland forest |  | White-sand ecosystem |  | Floodplain forest |  |
| --- | --- | --- | --- | --- | --- |
| Family | Species | Family | Species | Family | Species |
| Burseraceae | *Protium apiculatum* | Rubiaceae | *Ferdinandusa chlorantha* | Apocynaceae | *Malouetia tamaquarina* |
| Violaceae | *Rinorea guianensis* | Simaroubaceae | *Simaba guianensis* | Euphorbiaceae | *Hevea spruceana* |
| Euphorbiaceae | *Micrandropsis scleroxylon* | Primulaceae | *Cybianthus fulvopulverulentus* | Fabaceae | *Macrolobium acaciifolium* |
| Fabaceae | *Swartzia reticulata* | Fabaceae | *Aldina heterophylla* | Fabaceae | *Swartzia acuminata* |
| Humiriaceae | *Sacoglottis guianensis* | Humiriaceae | *Sacoglottis guianensis* | Humiriaceae | *Sacoglottis guianensis* |
| Lecythidaceae | *Corythophora alta* | Vochysiaceae | *Ruizterania retusa* | Lecythidaceae | *Couratari tenuicarpa* |
| Malvaceae | *Scleronema micranthum* | Malvaceae | *Scleronema micranthum* | Lecythidaceae | *Eschweilera tenuifolia* |
| Sapotaceae | *Ecclinusa guianensis* | Malvaceae | *Catostemma sclerophyllum* | Lauraceae | *Ocotea aciphylla* |
| Sapotaceae | *Manilkara elata* | Sapotaceae | *Manilkara bidentata* | Sapotaceae | *Manilkara bidentata* |
| Sapotaceae | *Pouteria anomala* | Sapotaceae | *Pradosia schomburgkiana* | Sapotaceae | *Pouteria pachyphylla* |

### Sampling design

The bark and fresh leaf spectral data were collected directly in the field from mature trees with over 10 cm DBH, with leaves reaching the canopy. We selected a minimum of 10 individuals of 10 abundant species from each ecosystem (upland, white-sand, and floodplain forest) according to an *a priori* inventory to create the local models on the ATTO site (see Table 1). The spectral collections were carried out by ecosystems at different times: in the floodplain forest (*igapó*) it was in October 2022, when the plots were not flooded; in the white-sand ecosystem it was in March 2023 and in the upland forest in August 2023, following the phenology of the trees to ensure that the leaves were in the mature leaf phase.

### Spectral data

Spectral collections of different tree tissues in the field were conducted using a portable high-resolution spectroradiometer ASD FieldSpec^®^ 4, equipped with the contact probe accessory for bark spectral measurements on trunks and the leaf clip accessory for leaf spectra collection, with a black background. This portable instrument facilitated rapid data acquisition, capturing information for 0.1 seconds per spectrum. The spectral region covered wavelengths from 350 to 2500 nm, encompassing the visible (VIS = 350–700 nm), near-infrared (NIR = 700–1300 nm), shortwave-IR (SWIR I = 1300–1900) nm, and shortwave-IR (SWIR II = 1900–2500 nm) range of the electromagnetic spectrum. The instrument offered a resolution of 3 nm to 700 nm and 10 nm to 2500 nm, measuring the sample’s reflectance and generating 2,151 data points.

The spectral signature pattern of the species has distinct curves between the three tissues evaluated: outer bark, inner bark, and fresh leaves (Fig. 2). To ensure the cleanliness of the instrument and prevent contamination with resins, latex, gum, sap, and dust, as well as to protect the lens of the contact probe, a glass slide (microscope slide) adapted to the optical reader was utilized to collect the trunk samples (outer and inner bark). Before each measurement, the equipment underwent calibration with Spectralon^®^ in conjunction with the glass slide. A minimum of three spectral readings of the outer bark and three spectral readings of the inner bark were recorded for each tree. The collection strategy involved obtaining spectra at different points around the tree to maximize the capture of individual variation (Hadlich *et al*., 2018). Spectra were collected from the most preserved part of the outer bark to ensure high-quality data. For inner bark collection, a machete delicately removes the outer bark until reaching the inner bark without passing through the cambium and the wood, i.e., without injuring the tree. The bark of the tree is formed by non-living outer bark and inner bark with living tissues, including all the tissues from the vascular cambium to the periderm, formed by three layers: suber, phellogen, and phelloderm (Esau & de Morretes, 1974; Ferri, 1999). This means the outer and inner bark have different spectral curve patterns (Hadlich *et al.,* 2018). Visual inspection of the spectra collected *in situ* guided the accuracy of the spectral curve of each bark tissue according to the method used by Hadlich *et al*. (2018). If the curve deviated from the standard for the tissue, especially in the inner bark, additional collections were made by scraping more of the tissue until the data fidelity was guaranteed (see an example of the bark spectral pattern in Fig. 2). Leaf spectral collections involved harvesting branches from the uppermost portions of the trees, which a professional climber facilitated. Approximately 20 mature, healthy leaves were selected from each individual, and readings were taken using four overlapping leaves on the adaxial side (Jacquemoud & Ustin, 2019). This approach yielded five spectral readings per individual. Calibration for each leaf measurement was accomplished using the white side of the leaf clip. These meticulous and standardized procedures ensured the acquisition of high-quality spectral data, representative of different tree tissues, forming the foundation for the subsequent analyses and model development.

**Fig. 2.**
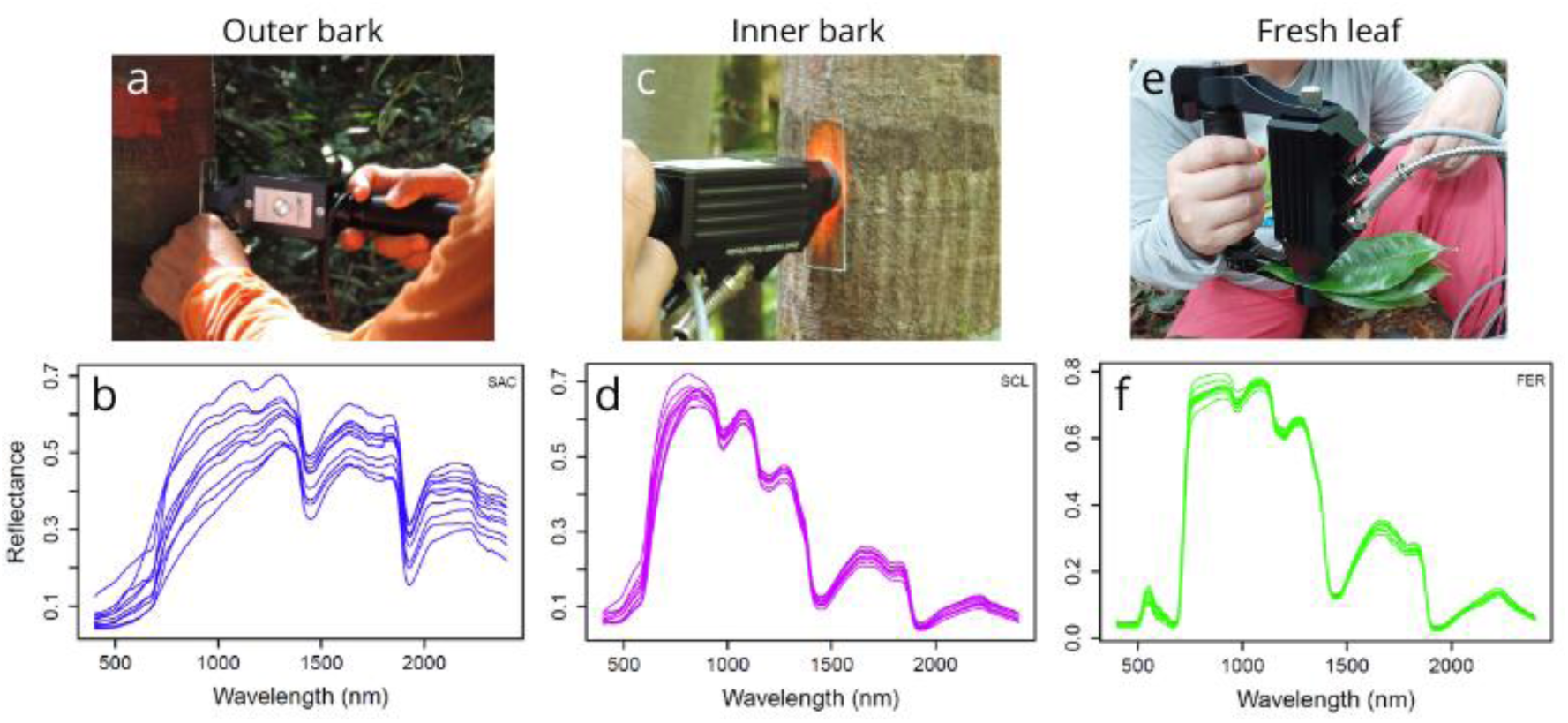
Spectral reflectance curve patterns for each tissue: panels a, c, and e show examples of field measurements for the outer bark; inner bark; and fresh leaf respectively; panels b, d, and f show the corresponding spectral curves.

### Data analysis

To evaluate the potential of the *in situ* collected spectra of the bark (inner and outer) and leaves for species recognition, spectral models were constructed for different tree tissues within each ecosystem. The initial phase involved preprocessing the spectral data to assess the quality and incorporated careful visualization of spectral curves and principal component analysis (PCA) to identify spectral behavior and outliers. Subsequently, wavelengths within 350–399 nm and 2401–2500 nm were excluded to eliminate equipment-induced noise.

Models based on Linear Discriminant Analysis (LDA) were then developed for the different tree tissues in each ecosystem to assess their predictive capacity for species recognition. In the LDA models, the dependent variables were the species (data known *a priori*), and the independent variables comprised the reflectance obtained at each wavelength from individual readings (Hair *et al*., 2009). The predictive efficiency of the models was tested using two cross-validation techniques: leave-one-out (LOO) and holdout with 70/30 (70/30) (Kohavi, 1995), which is consistent with the methodologies employed by Durgante *et al*. (2013) and Hadlich *et al*. (2018). As the spectral data is highly collinear, the stepwise method was used to determine the most important variables for discriminating between species and reduce the variables’ collinearity, using a maximum of 1/3 of the number of samples (William & Titus, 1988). The models were built using the mean of the spectral readings per individual. Four spectral models were established: individual models for each ecosystem (Upland, white-sand and floodplain forest) based on the data set for each tissue (outer bark, inner bark and fresh leaf) and a general model that included all the individuals from the three ecosystems for each tissue. All data processing and analysis were performed using the open-source statistical software R (v 4.4.1; R Core Team, 2024), employing custom-made R scripts and add-on packages. The analysis of variance was carried out in Python 3 with Jupyter Notebook as the primary environment. The Python libraries pandas, NumPy and matplotlib.

## Results

The variation of the spectra signals changes between plant tissues and ecosystems (Fig. 3 and Fig. S1). The outer and inner bark tissues showed a similar pattern of variation in the spectra of the species in all ecosystems. The lowest variation of both tissues was in the floodplain forest (Fig. S1.A) and the highest variation in the white-sand ecosystem, with the greatest difference in the VIS region, showing 111% (CV) in the inner bark and 54.6% (CV) in the outer bark. Inner bark showed very similar mean spectral curves of the species and CV in the upland and floodplain forest (Fig. 3 and S1.A). The fresh leaf spectral signal was the tissue with the lowest variation in all ecosystems (S1.B), with a maximum variation of 16.5 % (CV) in the upland forest, 23.9 % (CV) in the white-sand ecosystem and 29.9 % (CV) in the floodplain forest, in the VIS region. Overall, the region with the greatest variations was the VIS region, followed by SWIR II and then SWIR I and NIR.

**Fig. 3.**
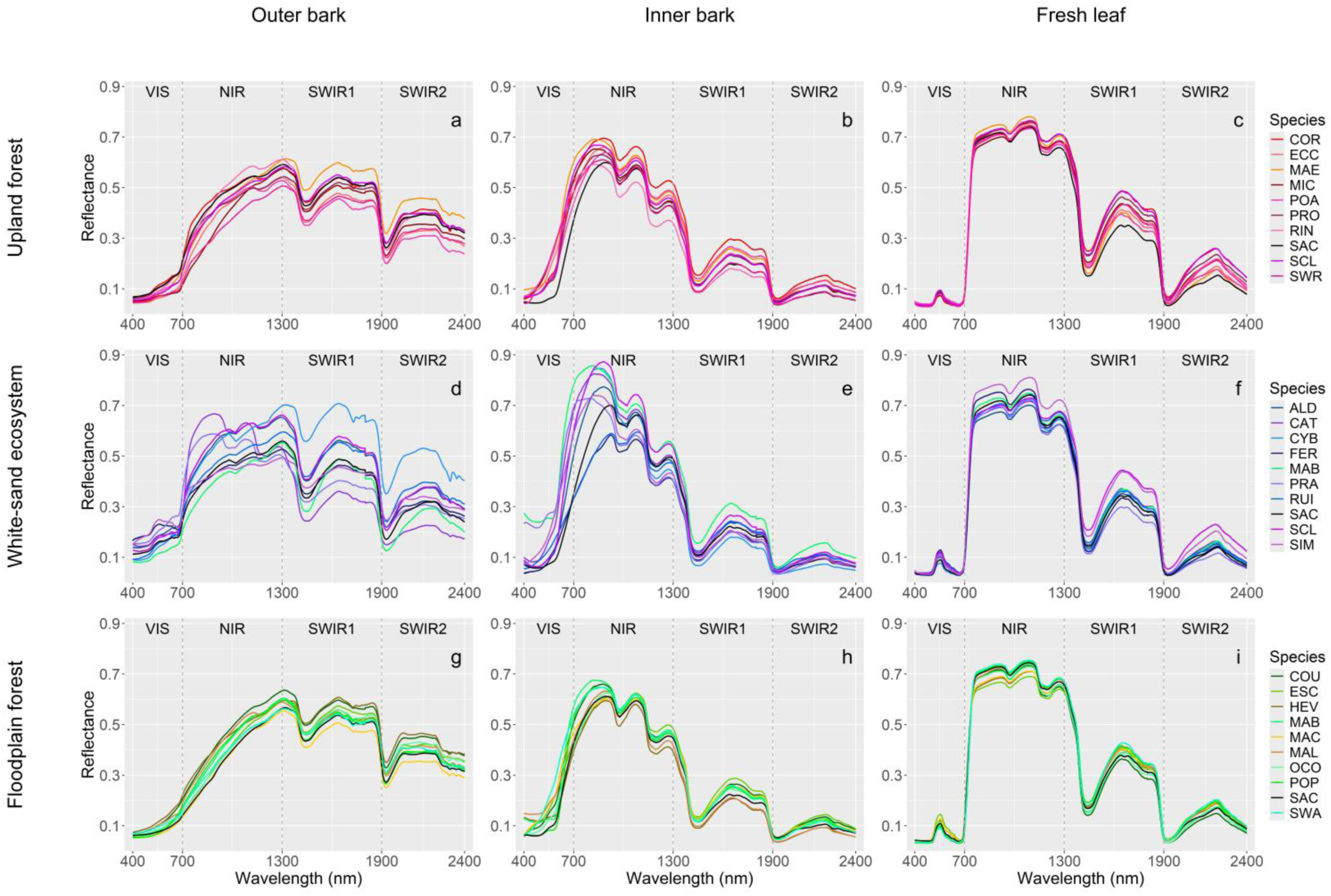
Mean spectral reflectance curves of tree species per tissue type across ecosystems. Species abbreviations are as follows: upland forest (COR = *Corythophora alta*, ECC = *Ecclinusa guianensis*, MAE = *Manilkara elata*, MIC = *Micrandropsis scleroxylon*, POA = *Pouteria anomala*, PRO = *Protium apiculatum*, RIN = *Rinorea guianensis*, SAC = *Sacoglottis guianensis*, SCL = *Scleronema micranthum*, and SWR = *Swartzia reticulata*); white-sand ecosystem (ALD = *Aldina heterophylla*, CAT = *Catostemma sclerophyllum*, CYB = *Cybianthus fulvopulverulentus*, FER = *Ferdinandusa chlorantha*, MAB = *Manilkara bidentata*, PRA = *Pradosia schomburgkiana*, RUI = *Ruizterania retusa*, SAC = *Sacoglottis guianensis*, SCL= *Scleronema micranthum*, and SIM = *Simaba guianensis*); floodplain forest (COU = *Couratari tenuicarpa*, ESC = *Eschweilera tenuifolia*, HEV = *Hevea spruceana*, MAB = *Manilkara bidentata,* MAC = *Macrolobium acaciifolium*, MAL = *Malouetia tamaquarina*, OCO = *Ocotea aciphylla*, POP = *Pouteria pachyphylla*, SAC = *Sacoglottis guianensis*, and SWA = *Swartzia acuminata*).

Species discrimination was successfully achieved using spectral signatures across all tested tree tissues and ecosystems. Taxonomic prediction accuracy ranged from 78.47% to 100%, depending on the validation method (LOO and 70/30 holdout) and using mean spectral readings per individual (see Table 2). Among the tissues, outer bark had the lowest prediction accuracy, ranging from 78.5 to 89%, while inner bark models achieved accuracies between 92.2% and 97%. Fresh leaf models exhibited the highest accuracy, varying between 92.4% to 100%. In the outer bark models, the lowest predictions in the 70/30 holdout validation occurred in white-sand trees (79.6%), while the highest was in the floodplain forest trees (86.5%). Inner bark models had the lowest prediction in floodplain forests (92.2%) and slightly higher accuracy in upland and white-sand ecosystems (95.5%). For fresh leaf spectral models, the lowest accuracy was observed in floodplain forests (92.4%), while white-sand trees showed the highest accuracy (99.9%).

**Table 2.** Results of the linear discriminant analysis (LDA) for the three tissue types within each ecosystem and a general model encompassing all ecosystems. Accuracy was assessed using the leave-one-out (LOO) and holdout (70/30 split) cross-validations. n represents the number of individual trees used to build the models.

| Model | Validation | n | Outer bark | Inner bark | Fresh leaf |
| --- | --- | --- | --- | --- | --- |
| Upland forest | LOO | 101 | 86% | 97% | 98% |
|  | 70/30 |  | 78.5% | 95.5% | 97.2% |
| White-sand ecosystem | LOO | 105 | 82% | 95% | 100% |
|  | 70/30 |  | 79.6% | 95.2% | 99.9% |
| Floodplain forest | LOO | 102 | 89% | 93% | 96% |
|  | 70/30 |  | 88.3% | 92.2% | 92.4% |
| General model | LOO | 308 | 86% | 97% | 98% |
|  | 70/30 |  | 86.5% | 96.2% | 97.7% |

Figure 4 presents confusion matrices highlighting misclassified species in each ecosystem based on the LOO validation. Correctly classified individuals are positioned along the diagonal, whereas misclassified trees appear outside the diagonal. In upland forests, the outer bark model misclassified 13 out of 101 individuals evaluated using the LOO validation. In contrast, the inner bark and fresh leaf models were misclassified by only three and two individuals, respectively. Notably, one misclassified individual was familiar across all three models belonging to *Pouteria anomala*. In the white-sand ecosystem, the outer bark misclassified 13 out of 105 individuals evaluated using the LOO validation, while the inner bark model misclassified only five trees. The fresh leaf model, however, attained 100% accuracy. In the floodplain forest, the outer bark model misclassified 9 out of 102 individual trees, performing the best model among forest types. The inner bark model misclassified seven individuals, while the fresh leaf model misclassified only four. In this ecosystem, the LOO validation (Fig. 4) revealed that *Hevea spruceana* and *Couratari tenuicarpa* were commonly misclassified across the outer bark, inner bark, and fresh leaf models.

**Fig. 4.**
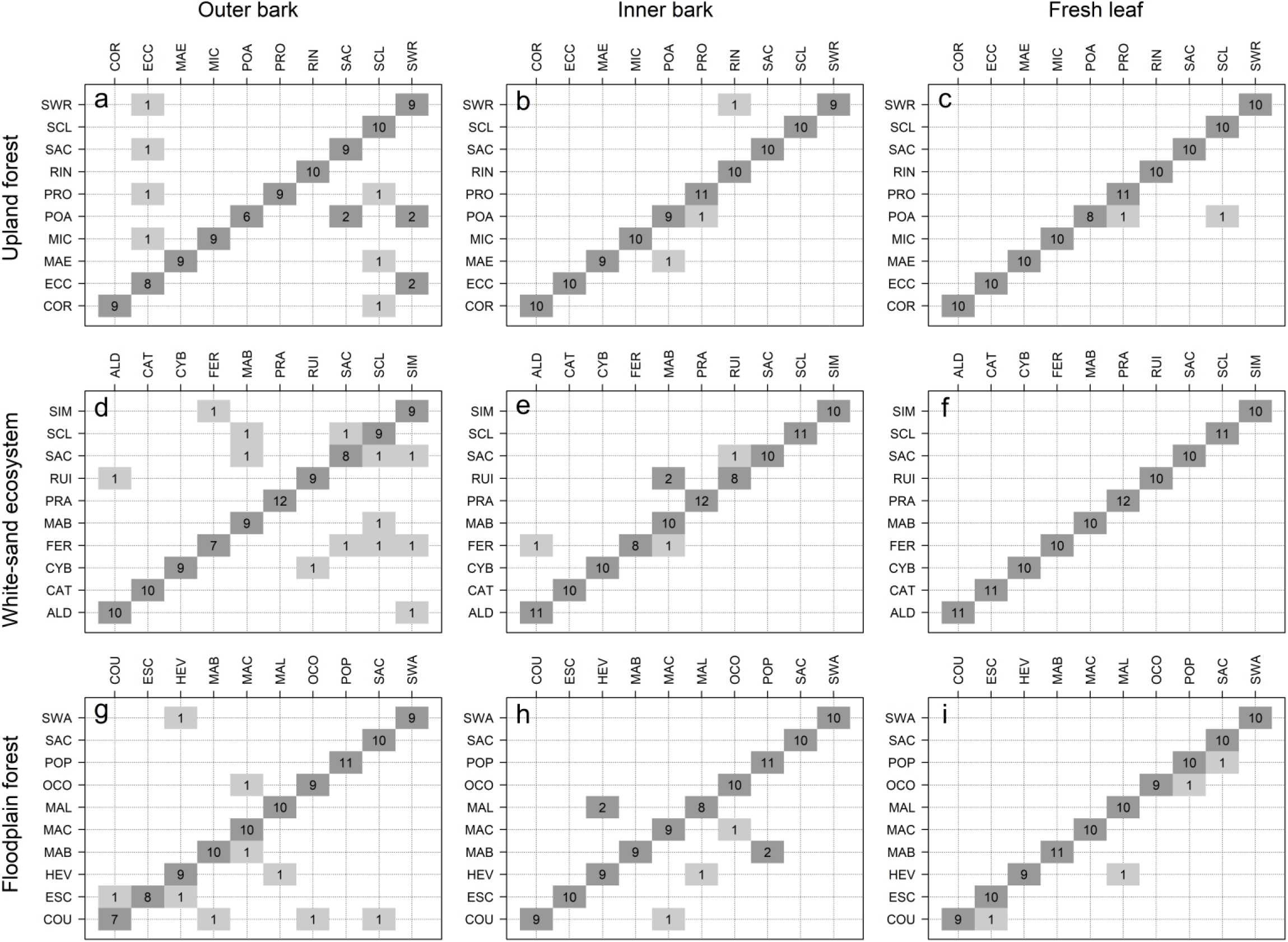
Confusion matrices resulting from LOO validation for each model tested for across different ecosystems. The diagonal represents correct predictions of species identification, while off the diagonal line are the incorrect predictions using the spectroscopic method. Species listed horizontally correspond to those identified by a taxonomist, while those listed vertically correspond to species “identified” using spectroscopy. Species abbreviations are as follows: Upland forest (COR = *Corythophora alta*, ECC = *Ecclinusa guianensis*, MAE = *Manilkara elata*, MIC = *Micrandropsis scleroxylon*, POA = *Pouteria anomala*, PRO = *Protium apiculatum*, RIN = *Rinorea guianensis*, SAC = *Sacoglottis guianensis*, SCL = *Scleronema micranthum*, and SWR = *Swartzia reticulata*); white-sand ecosystem (ALD = *Aldina heterophylla*, CAT = *Catostemma sclerophyllum*, CYB = *Cybianthus fulvopulverulentus*, FER = *Ferdinandusa chlorantha*, MAB = *Manilkara bidentata*, PRA = *Pradosia schomburgkiana*, RUI = *Ruizterania retusa*, SAC = *Sacoglottis guianensis*, SCL = *Scleronema micranthum*, and SIM = *Simaba guianensis*); floodplain forest (COU = *Couratari tenuicarpa*, ESC = *Eschweilera tenuifolia*, HEV = *Hevea spruceana*, MAB = *Manilkara bidentata,* MAC = *Macrolobium acaciifolium*, MAL = *Malouetia tamaquarina*, OCO = *Ocotea aciphylla*, POP = *Pouteria pachyphylla*, SAC = *Sacoglottis guianensis*, and SWA = *Swartzia acuminata*).

When combining all the individuals from the three ecosystems into a single general model, prediction accuracy remained high, ranging from 86% to 98% (Table 2). Fig 5 illustrates the misclassifications. The outer bark model had the lowest accuracy, with 26 misclassified individuals from 308. In contrast, the inner bark and fresh leaf models performed similarly, achieving 97% and 98% accuracy in the LOO validation, respectively, with only nine misclassified individuals in the inner bark and seven in the fresh leaf model.

**Fig. 5.**
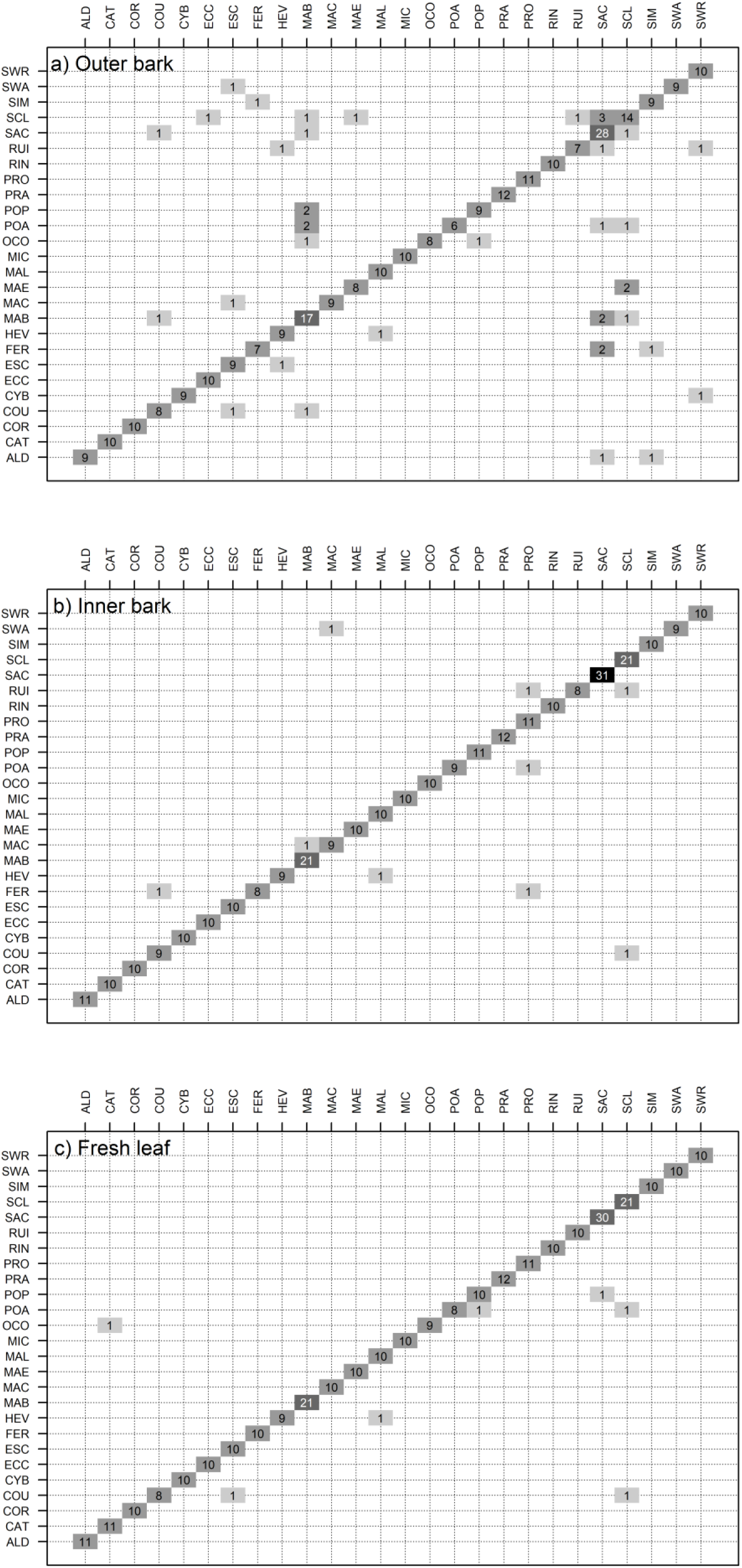
Confusion matrix resulting from LOO validation of the general model from fresh leaf. The diagonal line represents the correct predictions for species identification and those off the diagonal line are the incorrect predictions using the spectroscopic method. Species listed horizontally correspond to species identified by a taxonomist, while those listed vertically correspond to species “identified” using spectroscopy. *Species are the same as table 1 and Figs. 3 and 4.

Using the stepwise technique, the most informative characteristics for species classification were identified for each tissue type using the general models (Fig. 6). The outer bark model (fig 6.a) selected the 52 most informative variables, with the SWIR I region (1300 nm–1900 nm) being the most representative. The inner bark model (fig 6.b) relied on 66 selected variables, primarily from the VIS and SWIR I regions. For fresh leaves (fig 6.c), 78 variables were selected, with 41 in the VIS region. Table S1 presents all selected variables for each of the three tissue models.

**Fig. 6.**
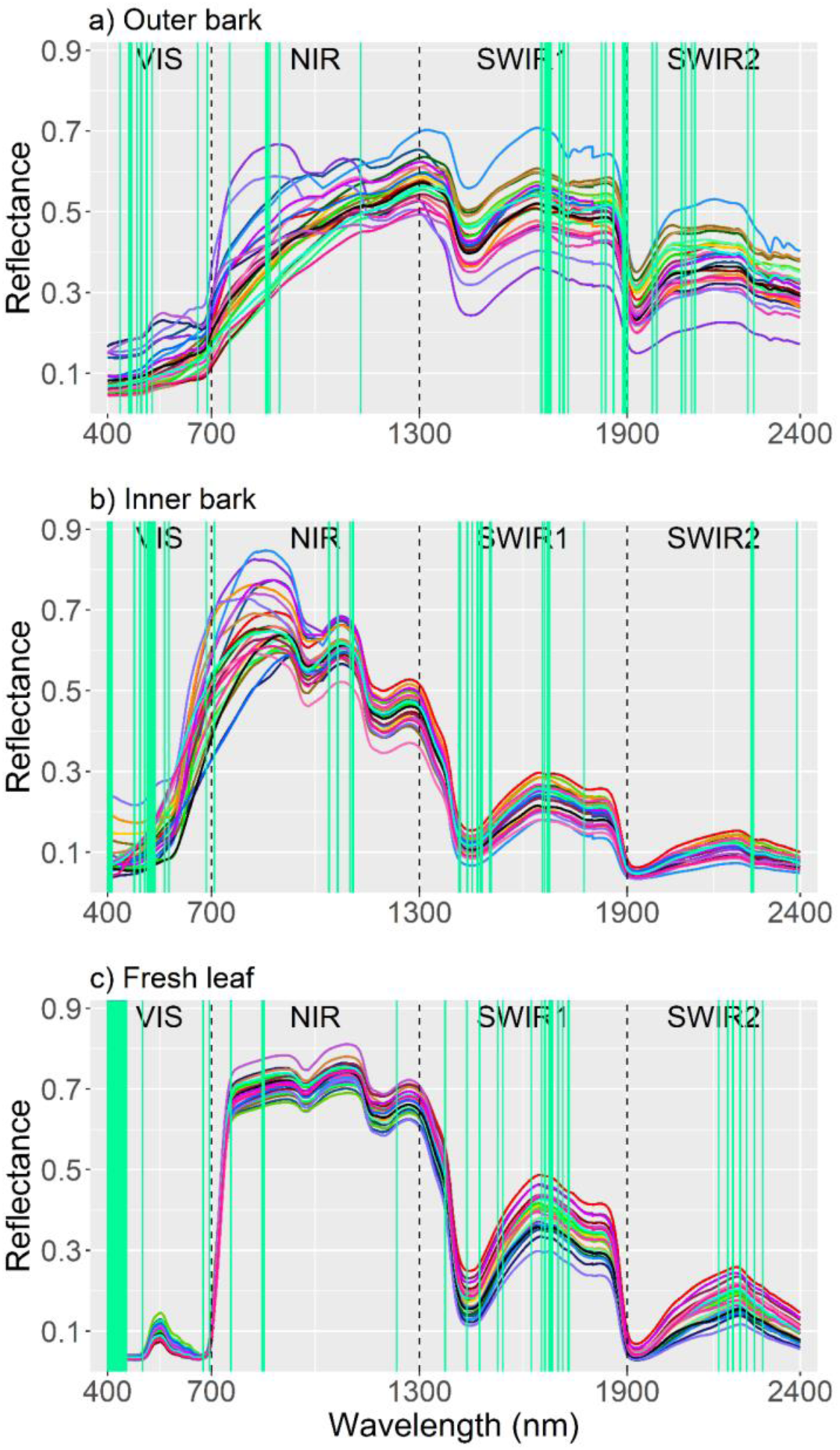
Mean spectral reflectance curves of all species sampled at the ATTO site, with the wavelengths selected (vertical green lines) for the general model by tissue type using the stepwise selection technique. *Species are the same as table 1 and figs 3, 4 and 5.

## Discussion

This study assessed the potential of spectral signatures from different tree tissues—outer bark, inner bark, and fresh leaves—for species identification across three distinct Amazonian ecosystems: upland, white-sand, and floodplain forest. Our results demonstrate that species discrimination was successfully achieved across all tested tissues and ecosystems. Fresh leaves exhibited the highest classification accuracy among the tissues analyzed, followed by inner and outer bark. Furthermore, we evaluated the feasibility of developing a generalized spectral model applicable across these three forest types. Despite the variations in spectral responses, the model achieved high accuracy in species recognition for the tested tissues. These findings highlight the promising potential of NIRS spectroscopy as a powerful tool for enhancing biodiversity inventories and advancing field-based species identification in the Amazon.

The spectral models based on fresh leaves showed the highest effectiveness for species recognition in the field for all ecosystems. The spectral reflectance of the leaves is determined by their chemical adaptations to environmental conditions, including climate, nutrient availability, phenology, age of the individual, lighting (Asner, 1998; Asner *et al*., 2014; Ponzoni, 2002), which influence their biochemical composition. The most accurate fresh leaf model was in the white-sand ecosystem, where the leaves exhibit scleromorphic traits, being thicker and enriched with secondary compounds to avoid herbivory, potentially enhancing their spectral signal. For the floodplain forest, the fresh leaf model showed slightly lower accuracies (92% and 96%) than those observed in the white-sand and upland. These results may have been influenced by leaf phenology, since in the field there is no such control, although only adult individuals and the preference for mature, healthy leaves were used. Our findings corroborate those of Prosper *et al*. (2014) and Liu *et al*. (2012) who successfully discriminated plant species *in situ* in wetlands with leaf spectra using different classification methods. However, collecting leaves requires climbing trees, an expensive and time-consuming field practice, especially considering the great heights of the trees in the Amazon. All the same, these findings are relevant for advancing hyperspectral and multispectral models using remote sensing technologies in biodiversity assessments (Feret & Asner, 2012; Cavender-Bares *et al*., 2020). In addition to species discrimination in hyperdiverse environments, leaf spectra can also be robustly applied in differentiating between invasive and native species, which is important for the management of conservation areas (Mallmann *et al.,* 2023).

The inner bark was the second-best tissue for species identification, with the spectral model accuracy decreasing by less than 5% compared to fresh leaves across all forest types. This slight reduction suggests that inner bark is a viable alternative for species classification in forest inventories, balancing efficiency and practicality. Outer bark, while less effective, still achieved an accuracy of approximately 89%, only 18% lower than fresh leaves. Although this reduction is notable, it remains a reasonable outcome given the challenges associated with outer bark spectroscopy. Our findings align with Hadlich *et al*. (2018), who demonstrated similar results for upland forest trees in the Central Amazon. Our study expands these insights across different forest types, reinforcing the potential of inner and outer bark spectroscopy for species identification in diverse Amazonian ecosystems.

The lower accuracy of outer bark models can be attributed to their exposure to environmental factors, including dust and microorganisms (e.g. lichens), and substantial variation in bark thickness between species (Paine *et al*., 2010; Rosell, 2016). These effects were captured with the spectral variation within the tissue between species. They were pronounced in the white-sand ecosystem, probably due to the scleromorphic characteristics of these forests, where trees are functionally adapted to water scarcity, leading to distinct chemical and structural traits that affect spectral discrimination, such as bark thickness and texture. The high variability in bark morphology and texture contributed to more significant spectral variability. This variation in the outer bark, along with differences between species, also changes throughout the tree’s ontogeny. However, further studies assessing the spectral responding of bark at different ontogenetic stages may provide valuable insights. The rough, cracked, fissured and fenestrated barks for example have the limitation of light scattering at the time of collection with the ASD Fieldspec 4 equipment in the field, leading to less consistent spectral curves to the outer bark than the more uniform patterns observed in fresh leaves and inner bark. Because it is a tissue that is protected by the outer bark, the inner bark, however, presents the most conserved biochemical characteristics and has less intraspecific variation (Hadlich *et al*., 2018). In the field, with the removal of the outer bark, the surface becomes regular, facilitating the accuracy of the spectral curve exposed from inner bark. However, Juola *et al*. (2023) demonstrated that the use of *in situ* imaging spectroscopy of the outer bark of standing trees, together with information on the texture of the trunk bark, can significantly improve the accuracy of spectral models of ten boreal species, with an accuracy of between 93% and 97.5%. Thus the combination of spectroscopy with morphological data is a promising alternative for the development of accurate applications for identifying tree species and supporting inventory (Juola *et al*., 2022, 2023). On the other hand, for the floodplain forest, the model with the outer bark performed better between upland and white-sand, showing the lowest intraspecific variation and with a spectral curve pattern similar to that of upland. This is remarkable, considering that trunks experience long periods of flooding, which could accelerate the decomposition of their tissues, affecting predictions.

The general model’s results were also promising, as accuracy remained high for fresh leaves, inner bark, and outer bark models. This suggests species identification in forest inventories does not require separate models for each forest type. By incorporating spectra from individuals across upland, white-sand, and floodplain forests, the general model captured a broader range of phenotypic variation, integrating more comprehensive physical and chemical information. However, regarding the applicability of this technique across different regions of the Amazon, further studies are needed to assess the potential for regionalizing this local model. Two key factors must be carefully evaluated when testing the regionalization of the model: i. the spectral response of species in other ecosystems not tested in this study; ii. species with identification challenges may involve species complexes or distinct allopatric species that have yet to be formally identified or described. Using such models represents a significant step forward in improving the efficiency of species identification for biodiversity research and monitoring, ultimately contributing to more effective conservation strategies in the Amazon.

Identifying the most informative wavelength for species discrimination across different tissues is essential for optimizing spectral models, especially considering variations in the range of the electromagnetic spectrum captured by the equipment, spectral resolution, and the influence of chemical signals on classification accuracy. The VIS (400–700 nm) and SWIR I (1300–1900 nm) regions were the most important for distinguishing species across all three tested tissues, with some variations among them. The fresh leaves models used 48 wavelengths in the VIS region, 18 in the SWIR I, and only 5 in the NIR region (700–1300 nm). Fresh leaf spectral responses are a function of leaf structure, water content, and the concentration of biochemicals (Curran *et al*., 1992). The discriminatory power in the VIS region was evidenced in the spectral range corresponding to the blue-green transition (400-499 nm) (Hennessy *et al*., 2020). In this region, the optical properties of the leaves remain stable due to the presence of biologically active pigments, such as chlorophyll and carotenoids (Gausman, 1982; Jones & Vaughan, 2010) that play a crucial role in this discrimination process. The NIR region, which reflects cell structure (Ponzoni *et al.,* 2015), had only 5 wavelengths selected by the spectral model. Additionally, specific wavelengths in the shortwave infrared (SWIR II) region (2.100–2300 nm) provide valuable information on cellulose and lignin content (Curran, 1989).

The spectral model of the outer bark selected 34 informative wavelengths in the SWIR I and II regions, of which 16 specific wavelengths belonged to lignin, hemicellulose, cellulose, aromatic compounds, and extractives (Jones *et al*., 2006; Workman & Weyer, 2008; Schwanninger *et al*., 2011) being the most representative region. For the inner bark, the spectral models selected 32 in the VIS region, and also detected 11 wavelengths belonging to lignin, cellulose, and water (Workman & Weyer, 2008), with VIS and SWIR I being the most important regions. When comparing the most informative wavelengths for species discrimination between outer and inner bark using the general model (upland, white-sand, and floodplain forests) to those presented by Hadlich *et al*. (2018) for upland forest, distinct differences emerged. In the present study, the most informative wavelengths for the outer bark were in SWIR I, whereas Hadlich *et al*. (2018) recognized VIS and NIR, as the most relevant. For inner bark, the selection of the most informative regions in this study were VIS and SWIR I, while Hadlich *et al*. (2018) identified VIS and NIR as the key regions. The selection of the most informative wavelength between models can be influenced by differences in the dataset, such as species composition and environmental variation. Juola *et al*. (2022) found selected features within the visible and NIR range for the outer bark. Therefore, the selection of variables that best discriminate between species must be refined because, in addition to the data set, this information can be influenced by different equipment with different resolutions and spectral range.

Spectroscopy is a valuable technique for improving tree biodiversity inventories in the field. However, some equipment-related limitations can be refined. The ASD FieldSpec 4 is an expensive device, making it difficult to popularize this technique among forest companies and scientific institutions. Additionally, the equipment weighs around 8 kg and requires a computer connection and battery power for operation, which makes it challenging to use in dense forest environments where accessing trees can be difficult. However, to make this technique more widely accessible to the field, we expect smaller, more affordable, and high-precision spectrometers, such as a small device from the ASD FieldSpec 4 or other spectral companies. The development and adoption of such equipment would help to expand the use of spectroscopy, facilitating the implementation of quality control in species identification and improving biodiversity inventories in tropical forest monitoring.

## Acknowledgements

This study was funded by the German - Brazilian project ATTO (Amazon Tall Tower Observatory) that was supported by German Federal Ministry of Education and Research (BMBF contract 01LK1602F and 01LK2101D) and the Brazilian Ministry of Science, Technology, Innovation, and Communication (FINEP/MCTIC, contract 01.11.01248.00). We also thank the PELD/MAUA project (process number 441811/2020-5 (CNPq), 01.02.016301.02630/2022-76 (FAPEAM); Chamada CNPq/MCTI/CONFAP-FAPs/PELD N° 21/2020); ATTO Hydrotraits project funded by the Conselho Nacional de Desenvolvimento Científico e Tecnológico (CNPq; Chamada N° 01/2022, process number 440166/2022-5); Universal Amazonas N° 006/2019 and Programa Mulher Faz Ciência N° 006/2024 (SPECTRA POP project) funded by the Fundação de Amparo à Pesquisa do Estado do Amazonas (FAPEAM). HLH received a doctoral scholarship from the Coordenação de Aperfeiçoamento de Pessoal de Nível Superior (CAPES) - Finance Code 001. CCV acknowledges the research grants awarded by CNPq (382803/2022-1 and #384501/2023-0) and FAPEAM (#01.02.016301.04984/2024-17). GBM was funded by the Gigante consortium NERC/NSF NE/Y003942/1; NSF-NERC DEB award #2241507. We would like to thank the MAUA group at the National Institute for Amazonian Research (INPA), the ATTO project, and PPG-BOT for providing scientific and logistical support. We also extend our thanks to all the field assistants and everyone involved in the logistic supportand.

## Author contributions

HLH, FMD, JS, FW planned and designed the research; HLH, CCV, CLM, MLGC, PAdeS, carried out the measurements and conducted fieldwork; CCV and LOD confirmed the taxonomic identification of the species; HLH, FMD, CCV, CLM performed statistical analyses; MTFP, FW, JS and LOD provided the inventory data; JS, FW and MTFP provided financial and logistical support; HLH and FMD wrote the first draft of the manuscript, and all authors contributed substantially to revisions.

