## Supplementary material for "Scaling up tree diversity inventories across Amazonian ecosystems using field spectroscopy": Fig S1 and Table S1

**New Phytologist Supporting Information**

**Fig. S1:** Graph of the coefficient of variation (CV) by tissue in each ecosystem on the left (A) and by ecosystem comparing the tissues on the right (B).

A

B

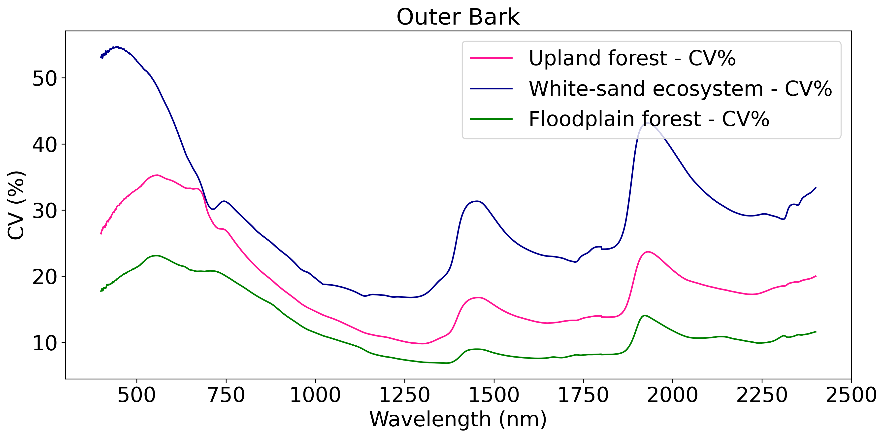

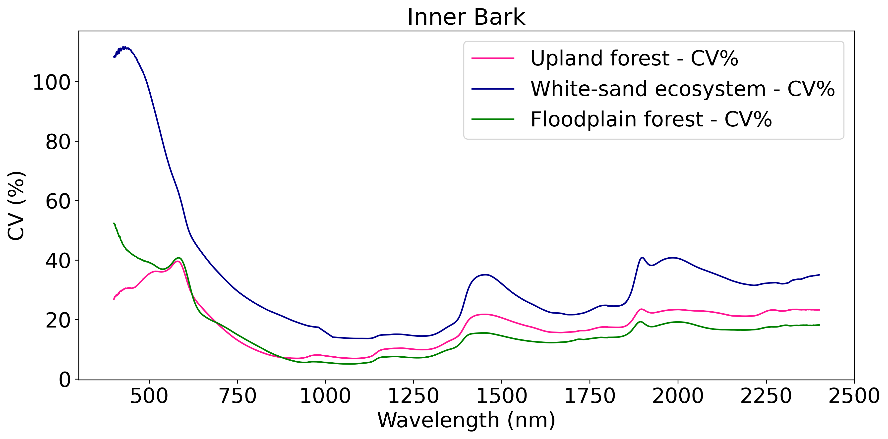

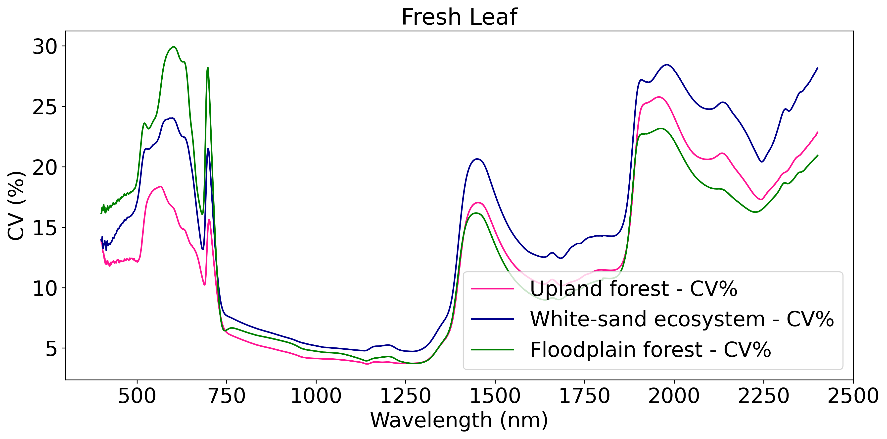

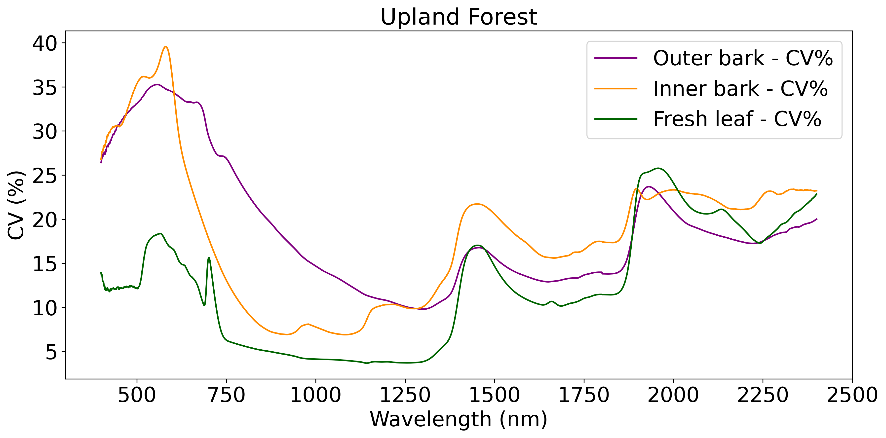

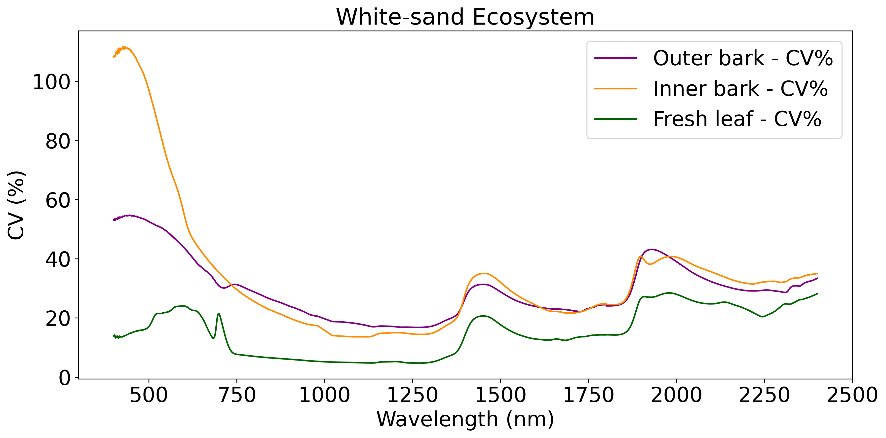

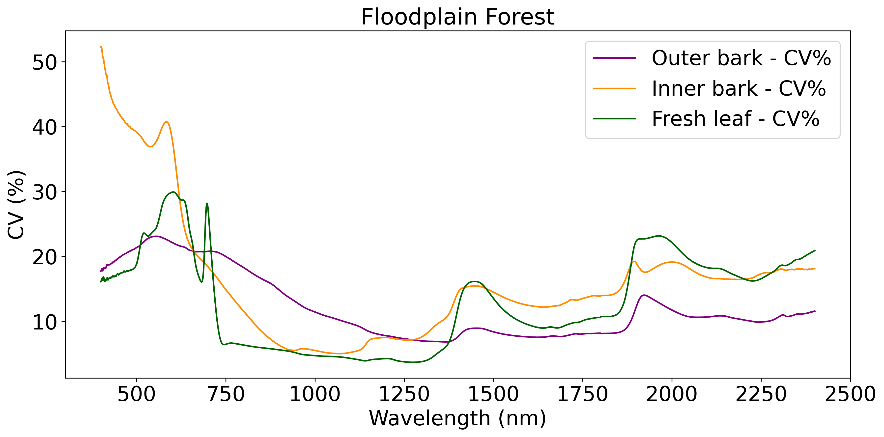

**Table S1:** Variables were selected using the stepwise method. Wavelengths (nm) of each tissue are shown by spectral region.

| **Spectral region** | **Outer bark** | **Inner bark** | **Fresh leaf** |
| --- | --- | --- | --- |
| **VIS** | 436 • 462 • 465 • 469 • 484 • 496 498 • 513 • 529 • 660 • 688 • 752 | 400 • 401 • 402 • 404 • 406 • 408 410 • 411 • 412 • 413 • 476 • 477 478 • 493 • 495 • 507 • 515 • 517 518 • 521 • 523 • 524 • 528 • 529 530 • 533 • 534 • 537 • 564 • 577 685 • 708 | 400 • 401 • 402• 403 • 404 • 406 407 • 408 • 409 • 410 • 411 • 412 413 • 414 • 415 • 416 • 417 • 418 419 • 420 • 421 • 422 • 424 • 425 426 • 427 • 428 • 431 • 433 • 434 435 • 436 • 438 • 439 • 440 • 441 443 • 444 • 445 • 447 • 448 |
| **NIR** | 856 • 859 • 862 • 868 • 896 • 1131 | 1038 • 1039 • 1040 • 1063 • 1064 1066 • 1099 • 1107 • 1109 • 1110 | 846 • 851 • 852 • 1235 • 1375 |
| **SWIR I** | 1651 • 1654 • 1663 • 1668 • 1672 1673 • 1674 • 1675 1676 • 1677 1680 • 1705 • 1714 • 1716 • 1717 1718 • 1729 • 1826 • 1838 • 1861 1862 • 1887 • 1890 • 1894 • 1897 | 1415 • 1417 • 1419 • 1438 • 1439 1452 • 1467 • 1468 • 1476 • 1480 1481 • 1504 • 1508 • 1655 • 1656 1662 • 1671 • 1672 • 1675 • 1676 1776 | 1437 • 1438 • 1474 • 1526 • 1541 1624 • 1653 • 1661 • 1663 • 1674 1679 • 1681 • 1685 • 1701 • 1714 1730 • 1731 • 1732 |
| **SWIR II** | 1903 • 1973 • 1986 • 2058 • 2068 2086 • 2096 • 2249 • 2266 | 2259 • 2265 • 2391 | 2165 • 2191 • 2206 • 2227 • 2245 2268 • 2292 |
